# Management factors influence *Salmonella* persistence in reused poultry litter over three successive flocks

**DOI:** 10.1101/2023.03.03.531069

**Authors:** Reed Woyda, Adelumola Oladeinde, Dinku Endale, Timothy Strickland, Jodie Plumblee Lawrence, Zaid Abdo

## Abstract

*Salmonella* infections are a leading cause of bacterial food-borne illness worldwide. Infections are highly associated with the consumption of contaminated food, and in particular, chicken meat. Understanding how management practices and environmental factors influence *Salmonella* populations in broiler chicken production may aid in reducing the risk of food-borne illness in humans. Utilizing whole genome sequencing with antimicrobial and heavy metal resistance, virulence factor and plasmid identification, we have characterized the genetic diversity of *Salmonella enterica* isolates (n = 55) obtained from broiler chicken litter. *S. enterica* isolates were recovered from the litter of broiler chickens over three consecutive flocks in four broiler houses on a single integrated farm in Georgia, USA. The chickens were raised under a newly adopted “No Antibiotics Ever” program and copper sulfate was administered via drinking water. *In-silico* serovar prediction identified three *S. enterica* serovars: Enteritidis (n = 12), Kentucky (n = 40) and Senftenberg (n = 3). Antimicrobial susceptibility testing revealed that only one *S*. Kentucky isolate was resistant to streptomycin, while the remaining isolates were susceptible to all antibiotics tested. Metal resistance operons, including copper and silver, were identified chromosomally and on plasmids in serovar Senftenberg and Kentucky isolates, respectively. Serovar Kentucky isolates harboring metal resistance operons were the only *Salmonella* isolates recovered from the litter of third flock cohort. These results suggest the addition of copper sulfate to drinking water may have selected for *S.* Kentucky isolates harboring plasmid-borne copper resistance genes and may explain their persistence in litter from flock to flock.

**Importance:** *Salmonella* foodborne illnesses are the leading cause of hospitalizations and deaths, resulting in a high economic burden on the healthcare system. Globally, chicken meat is one of the highest consumed meats and is a predominant source of foodborne illness. The severity of *Salmonella* infections depends on the presence of antimicrobial resistance genes and virulence factors. While there are many studies which have investigated *Salmonella* strains isolated from post-harvest chicken samples, there is a gap in our understanding of the prevalence and persistence of *Salmonella* in pre-harvest and in particular their makeup of antibiotic resistance genes, virulence factors and metal resistance genes. The objective of this study was to determine how on-farm management practices and environmental factors influence *Salmonella* persistence, as well as the antimicrobial resistance genes and virulence factors they harbor. In this study we demonstrate that broiler chickens raised without antibiotics are less likely to harbor antibiotic resistance, however the practice of adding acidified copper sulfate to drinking water may select for strains carrying metal resistant genes.

## Introduction

*Salmonella* is responsible for an estimated 1.2 million illnesses annually in the United States, 1 million of which are attributed to the consumption of contaminated food (Scallan *et al*., 2011). Chicken meat is the most consumed meat in the United States (USDA, Economic Research Service, 2022) and is a predominant source of food-borne illness (ADAS, 2016). *Salmonella* food-borne outbreaks are strongly associated with consumption of chicken meat (ADAS, 2016; Antunes *et al*., 2016). Moreover, the annual economic loss due to *Salmonella* infections is an estimated $11 billion in the United States (Wernicki, Nowaczek and Urban-Chmiel, 2017).

*Salmonella enterica* encompasses the most pathogenic species and consists of thousands of different serovars, some of which are host-specific while some maintain the ability to infect a broad range of hosts (Mezal *et al*., 2014). The United States Department of Agriculture Food Safety and Inspection Service surveillance fiscal year 2022 reports indicate the top *Salmonella* serovars isolates from domestic chicken samples are, in decreasing order, Infantis, Kentucky, Enteritidis and Typhimurium (USDA, 2022). In 2018, the Foodborne Diseases Active Surveillance Network (FoodNet) identified Enteritidis, Newport, and Typhimurium serovars were the most frequently identified as causing illness in humans (Tack et al., 2019). While some serovars are capable of infecting both chickens and humans (e.g., Enteritidis and Typhimurium), serovars such as Kentucky and Sofia are highly prevalent in poultry but have a low association with human outbreaks (Ferrari *et al*., 2019). Thus, the threat of *Salmonella* outbreaks in humans is serovar dependent, however the severity of disease and the efficacy of treatment depends on the strain’s virulence factor (VF) and antimicrobial resistance gene (ARG) repertoire. Importantly, VFs and ARGs may be shared across *Salmonella* serovars and, in some instances, across different bacterial genera if these genetic elements are located on specific plasmids or other mobile genetic elements (Fricke *et al*., 2009; Oladeinde *et al*., 2019).

*Salmonella* pathogenicity is dependent on the VF repertoire and pathogenesis proceeds through attachment to host tissues, invasion of host tissues, macrophage survival, replication and subsequent dissemination (Gao, Wang and Ogunremi, 2020). Virulence factors in *Salmonella* can be accumulated and, in combination, allow for successful host colonization and bypassing of host defenses. These virulence factors are typically encoded within *Salmonella* pathogenicity islands or on plasmids (Jajere, 2019; Gao, Wang and Ogunremi, 2020).

Antimicrobial therapy is the first choice treatment for *Salmonella* infections. However, the global rise and threat of antimicrobial resistance, due to the misuse of antibiotics in food-producing animals and human medicine, has put pressure on the broiler production industry to reduce usage of antimicrobials (United States Food and Drug Administration, Center for Veterinary Medicine, 2022). Difficulties in treating *Salmonella* infections occur when strains are resistant to multiple antimicrobials from distinct antimicrobial classes; such strains have been identified in poultry, beef and pork products (Gieraltowski *et al*., 2016; Bearson *et al*., 2019; Feng *et al*., 2020). Importantly, strains harboring ARGs which confer resistance to antimicrobials critically important to human medicine (World Health Organization, 2019), as well as those possessing VF increasing pathogenicity to humans, will pose a threat and will be harder to treat in the event contaminated food products reach consumers (Fricke *et al*., 2009; Jajere, 2019; Zakaria *et al*., 2022). Additionally, the use of heavy metals in chicken production, such as copper for growth promotion and/or sanitation of water lines, has led to the discovery that genes conferring metal resistance are often carried on the same mobile elements as ARGs and VFs (Dziewit *et al*., 2015; Pal *et al*., 2015; Bukowski *et al*., 2019; Souza *et al*., 2022). Resistance to metals may increase the virulence of a bacterial pathogen, e.g., overcoming iron toxicity and invading and colonizing different tissue types (Nairz *et al*., 2015), as well as give a competitive advantage to strains in production environments where metals are used for antimicrobial agents (Bearson *et al*., 2019). Thus, it is critical to understand how on-farm management practices and environmental factors influence strains harboring both ARGs, VFs and metal resistance genes to develop prediction, or management, methods to limit their occurrence in post-harvest chicken production.

Broiler chickens are raised on litter for an entire grow-out (0 – 56 days). Broiler litter is a complex matrix consisting of a decomposing plant-based bedding material, chicken excreta, feathers, and feed. Broiler chickens are copraphagic by nature and therefore will start to consume the litter upon placement in a broiler house. Thus, it is reasonable to assume that chicks will be exposed to pathogens during both the initial stages of growth and throughout their grow-out through normal activities such as pecking and bathing. Many studies have reported the occurrence of *Salmonella* in broiler litter (Jones *et al*., 1991; Kelley *et al*., 1995, p. 19; Pope and Cherry, 2000; Line, 2002; Brooks *et al*., 2010; Shepherd *et al*., 2010; Velasquez *et al*., 2018), but only a few studies have examined the genetic factors that allow *Salmonella* to persist in pre-harvest broiler production (Roberts *et al*., 2013; Vaz *et al*., 2017; Shang, Wei and Kang, 2018; Dunn *et al*., 2022).

The objective of this study was to determine how management practices and environmental factors affect the persistence of *Salmonella*, as well as their VFs, ARGs and metal resistance genes, in pre-harvest broiler production. In this study, we performed an in-depth genomic characterization of *Salmonella* isolates recovered from the litter of four co-located broiler chicken houses over three consecutive flocks (Oladeinde *et al*., 2023). We previously reported that these isolates were unequally distributed across the litter of the broiler houses and showed that the probability of detecting *Salmonella* was higher for the first broiler cohort raised on litter compared to cohorts 2 and 3 (Oladeinde *et al*., 2023). Here, we performed antimicrobial susceptibility testing and whole genome sequencing on 55 isolates recovered from the litter (Oladeinde *et al*., 2023). While antibiotic susceptibility testing resulted in one isolate displaying phenotypic resistance, our genomic characterization identified a potential beta-lactamase gene as well as copper and silver resistance operons. Characterization of the antimicrobial resistance genes, biocide and metal resistance genes and virulence factors revealed a large core set of genes with differences explained by isolates’ serovar. Only isolates harboring specific copper and silver resistance operons were detected over the three consecutive flock cycles. Together these results suggest the on-farm management practice of adding copper sulfate to the drinking water, commonly practiced in the industry, may provide means for *Salmonella* persistence over multiple flock cohorts.

## Results

### *Salmonella* serovar occurrence differed across grow-out periods, flocks, and broiler houses

*Salmonella* prevalence in litter collected from four broiler house floors was determined by direct and selective enrichment plating of litter eluate on both Brilliant Green Sulfur (BGS) and Xylose Lysine Tergitol-4 (XLT-4) agar (Oladeinde *et al*., 2023). *Salmonella* was detected in 10.76% (31/288) of litter samples over 3 consecutive flock cohorts across 4 co-located broiler houses (Oladeinde *et al*., 2023). Fifty-five *Salmonella* isolates were selected for whole genome sequencing. These isolates were obtained either through direct plating or enrichment methods and encompassed all unique serogroups (Oladeinde *et al*., 2023). *In silico* serovar prediction identified three different serovars: *S.* Enteritidis (n = 12), *S.* Kentucky (n = 40), and *S.* Senftenberg (n = 3) (**Table S1**). A majority of isolates (85.5%, n = 47) were recovered during the late grow-out (32-38 days old chickens). Each serovar identified was present in both early (4-14 days old chickens) and late grow-out phases (**Table S1**). All *S.* Enteritidis isolates were identified as ST11, all *S.* Kentucky isolates as ST152 (excluding SK34 and SK36 due to low sequencing coverage (< 20X)), and *S.* Senftenberg isolates were identified as ST14 (excluding SS26 due to low sequencing coverage (< 20X)). Multi-locus sequence typing (MLST) was performed using the PubMLST website and the MLST software (Jolley and Maiden, 2010).

*Salmonella* occurrence differed over multiple flock cohorts (Table 1, (Oladeinde *et al*., 2023)) The 37 *Salmonella* isolates from flock 1 consisted of the following serovars: Enteritidis (n=9), Kentucky (n=25), Senftenberg (n=3) and a single unidentified isolate. The second flock cohort included 10 *Salmonella* isolates classified as Enteritidis (n=3) and Kentucky (n=7). The final flock cohort, flock 3, harbored only *S.* Kentucky isolates (n=8). The distribution of isolates and serovars also differed across the broiler houses (**Figure 1**). Houses 2 (33%, n = 18) and 3 (51%, n = 28) harbored majority of the sequenced isolates and the remaining 9 (16%) originated from house 4. *Salmonella* was not detected from any litter samples from house 1. *Salmonella* isolates sequenced from house 4 included a single *S.* Enteritidis isolate as well as a set of 8 *S.* Kentucky isolates which were all clustered together within the back section of the broiler house (**Figure 1**).

**Table 1.** Occurrence of Salmonella species in peanut hull litter over 3 grow-out cycles across 4 co-located broiler houses.

| House | Flock cohort 1 |  |  | Flock cohort 2 |  |  | Flock cohort 3 |  |  |
| --- | --- | --- | --- | --- | --- | --- | --- | --- | --- |
|  | S. Enteritidis | S. Kentucky | S. Senftenberg | S. Enteritidis | S. Kentucky | S. Senftenberg | S. Enteritidis | S. Kentucky | S. Senftenberg |
| 1 | 0 | 0 | 0 | 0 | 0 | 0 | 0 | 0 | 0 |
| 2 | 8 | 8 | 0 | 0 | 2 | 0 | 0 | 0 | 0 |
| 3 | 1 | 17 | 3 | 2 | 5 | 0 | 0 | 8 | 0 |
| 4 | 0 | 0 | 0 | 1 | 0 | 0 | 0 | 0 | 0 |

**Figure 1.**
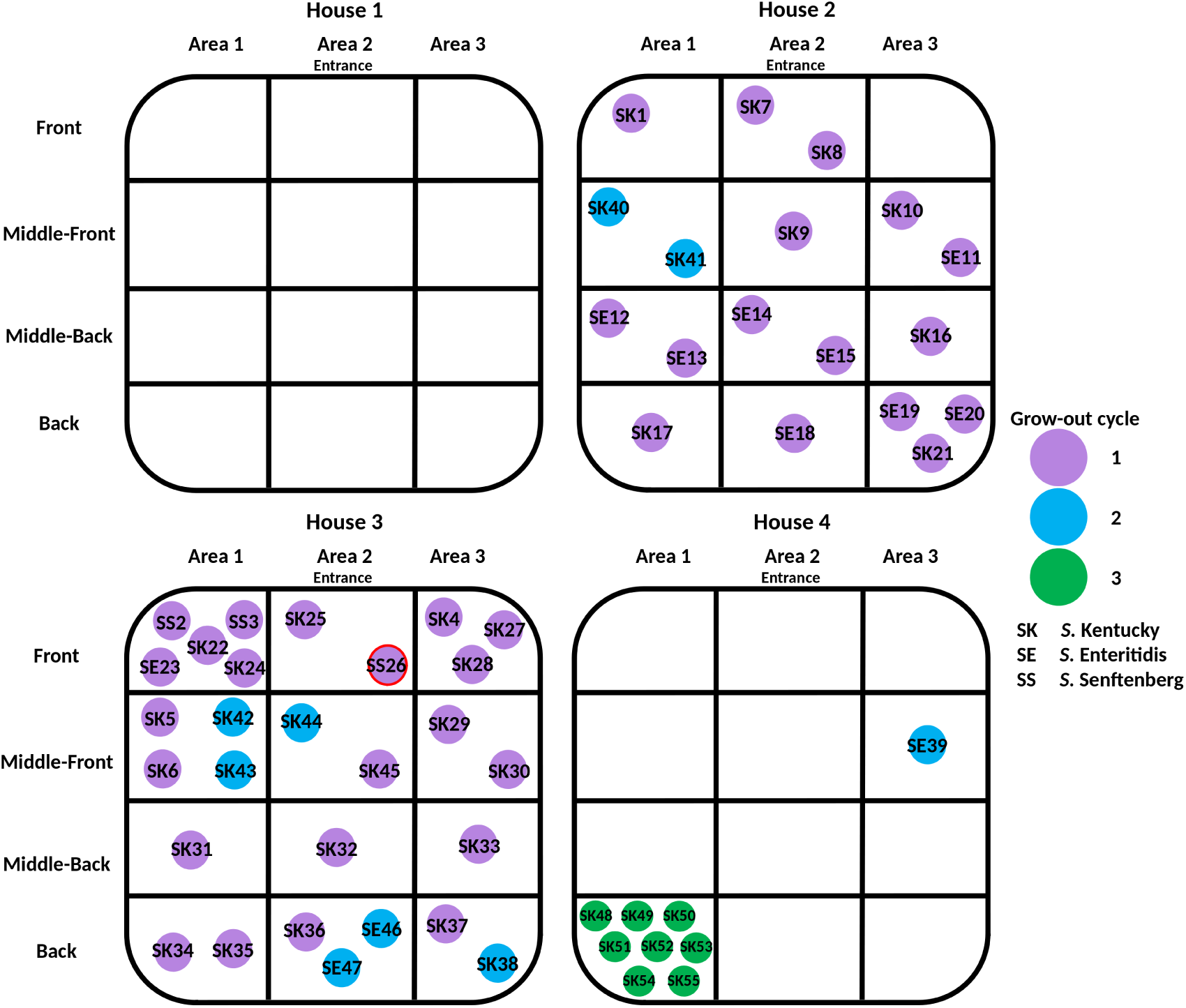
Visual representation of *Salmonella* species isolated from peanut hull litter within each section of each grow house. Samples were taken from 4 co-located grow houses over 3 grow-out cycles. Circles represent individual isolates labeled by their prospective serovar: *Salmonella* Kentucky (SK), *Salmonella* Enteritidis (SE) and *Salmonella* Senftenberg (SS). Circle color indicates which grow-out cycle an isolate was obtained from: grow-out cycle 1 (purple), grow-out cycle 2 (blue) and grow-out cycle 3 (green). The isolate outline in red was not included in further analysis due to poor sequencing coverage (SE23).

### Serovar Kentucky and Senftenberg harbored additional copper and silver resistance operons

Antibacterial biocides and metal resistance genes were identified using the experimentally confirmed resistance function BacMet database (Pal *et al*., 2014). A total of 125 genes with >80% identity to the reference database entry were identified and collectively conferred resistance to 74 unique compounds (**Table S2**). 108 of these genes were found to be present in >96.4% of isolates. Many copper and other metal homeostasis related genes were present in >96.4% of isolates (e.g., *copA* (detoxification of cytoplasmic Cu[I]) (Rensing and Grass, 2003), *cueP* (periplasm copper binding) (Pezza *et al*., 2016), and *zraP/zraR* (involved in intrinsic antibiotic and zinc resistance) (Rome *et al*., 2018)) (**Table S2**). In particular, the *silABCPRS* and *pcoABCDESR* resistance operons (silver and copper, respectively) were identified only in serovar Kentucky and Senftenberg isolates. (**Figure 2**). These operons are often clustered together on a plasmid and compose the copper homeostasis and silver resistance island (CHARSI) (Staehlin *et al*., 2016). While these operons appear to be chromosomally encoded within the Senftenberg isolates (contigs >390kbp), within Kentucky isolates these copper and silver operons appear to reside on an IncI1 plasmid.

**Figure 2.**
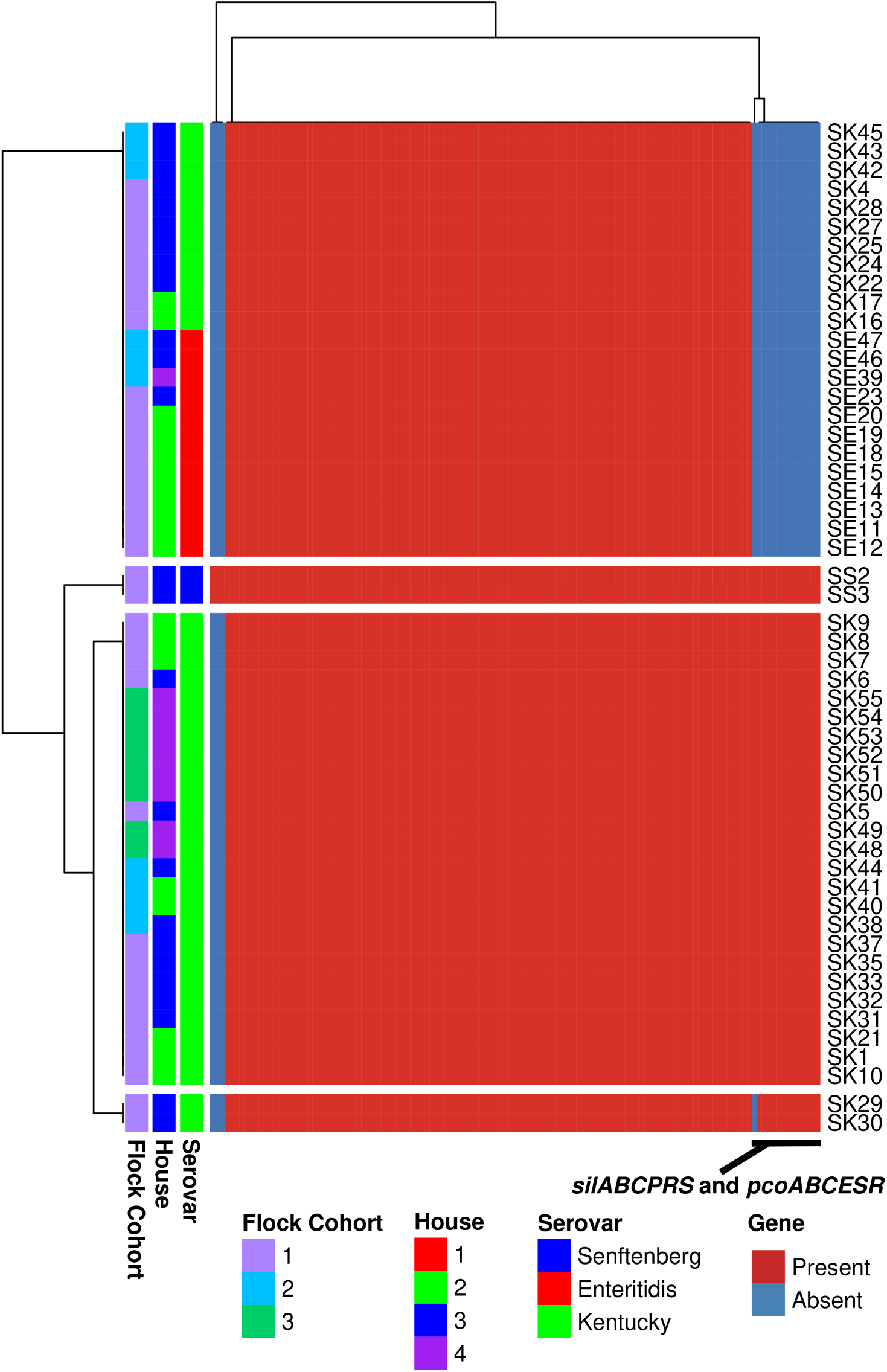
Heatmap of biocide and metal resistance genes. Genes conferring resistance to biocide and metal resistance were obtained via BacMet. Heatmap was generated in R v4.0.4 with pheatmap v1.0.12 (clustering_method = “average” (UPGMA)) using the filtered, >= 80% identity, presence/absence table of biocide and metal resistance genes (**Table S5**).

### *Salmonella* isolates share a majority of identified virulence factors

A total of 119 VFs were identified utilizing the Virulence Factor Database (VFDB) via ABRICATE and 86 (72%) VFs were present in all isolates (**Table S1**). Of the remaining 33 VFs, 19 were present in the majority of isolates (observed within > 70% of those isolates but less than 100%) while the remaining 14 where only observed in 4% - 21% of all isolates. Hierarchical clustering revealed that differences in VF across all isolates could be explained by an isolate’s serovar (**Figure 3**). The 119 VFs are involved in 12 different functions such as secretion systems (63%), fimbrial adherence determinants (20%), iron uptake (4%), adherence (2.5%) and magnesium uptake (1.7%) (**Table 3**, **Table S3**). Taken together, the *Salmonella* isolates harbored a similar core set of virulence factors and differences observed were explained by the isolates’ serovar.

**Figure 3.**
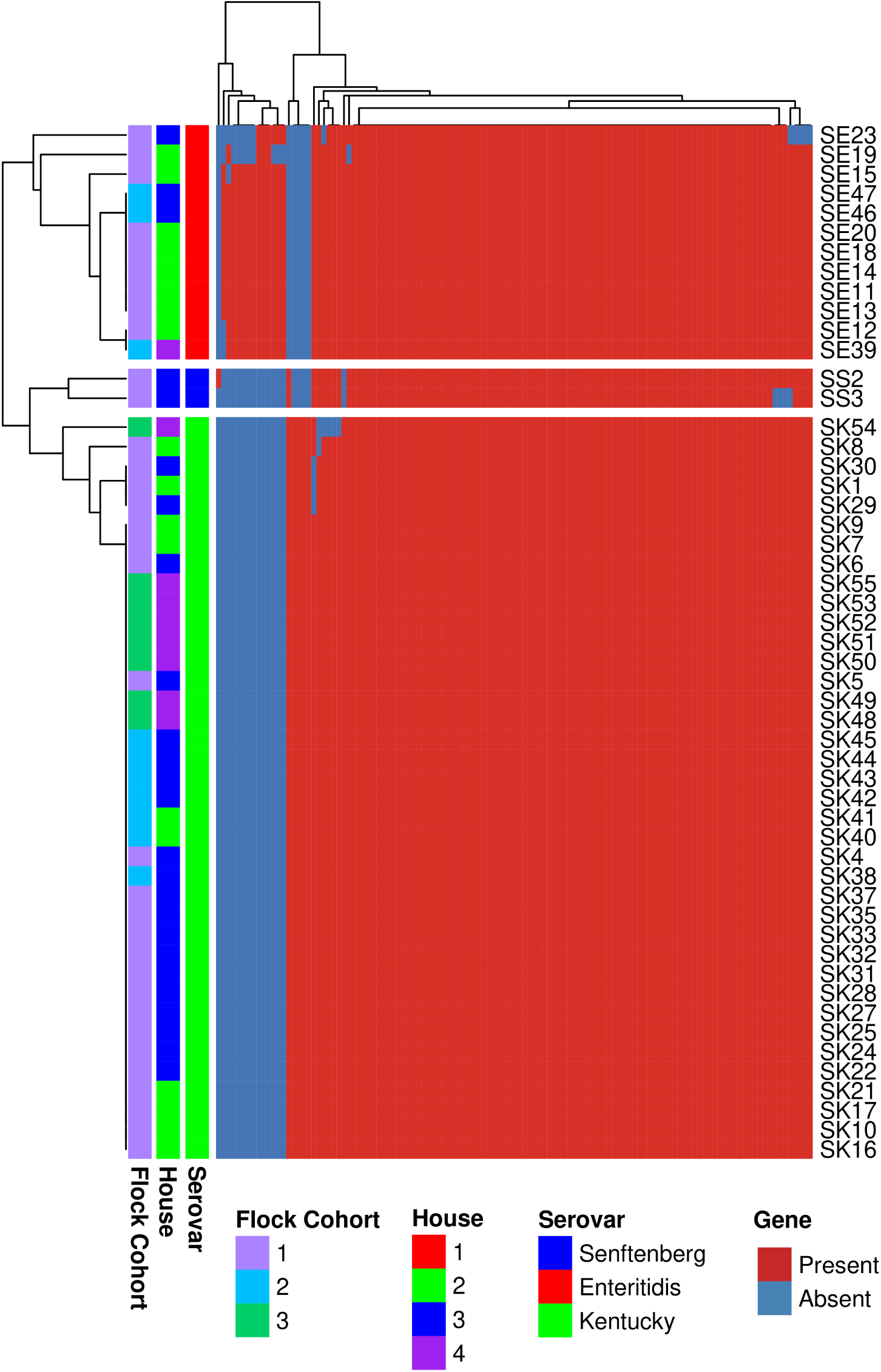
Heatmap of virulence factor genes. Virulence factors were identified from the Virulence Factor Database via ABRICATE. Heatmap was generated in R v4.0.4 with pheatmap v1.0.12 (clustering_method = “average” (UPGMA)) using the virulence factor data from **Table S1**.

**Table 2.** Enumeration of virulence factor functional categories of virulence factors identified from the Virulence Factor Database (VFDB) (via ABRICATE (Reads2Resistome)).

| Function | Number of genes |
| --- | --- |
| Secretion system | 75 |
| Fimbrial adherence determinants | 24 |
| Iron uptake | 5 |
| Nonfimbrial adherence determinants | 4 |
| Adherence | 3 |
| Magnesium uptake | 2 |
| Invasion | 1 |
| Macrophage inducible genes | 1 |
| Regulation | 1 |
| Serum resistance | 1 |
| Stress adaptation | 1 |
| Toxin | 1 |

**Table 3.** Virulence factors, antimicrobial resistance genes and plasmid replicons unique to specific serovars.

| Function | Subfunction | Gene/replicon<br>Name | Number of<br>Isolates | Serovar |
| --- | --- | --- | --- | --- |
| TEM beta-lactamase | extended-spectrum<br>beta-lactamase | *TEM-60* | 41 | Kentucky<br>and<br>Senftenberg |
| Plasmid replicon | Plasmid replicon | IncI1_1_Alpha | 39 | Kentucky |
| Plasmid replicon | Plasmid replicon | IncX1_3 | 39 | Kentucky |
| Plasmid replicon | Plasmid replicon | IncFIB.S._1 | 10 | Enteritidis |
| Plasmid replicon | Plasmid replicon | IncFII.S._1 | 8 | Enteritidis |
| Plasmid replicon | Plasmid replicon | Col156_1 | 1 | Kentucky |
| Plasmid replicon | Plasmid replicon | Col.MG828._1 | 1 | Kentucky |
| Secretion system | TTSS-2 translocated<br>effectors | sseK2 | 2 | Senftenberg |
| Secretion system | TTSS-2 translocated<br>effectors | sseI/srfH | 8 | Enteritidis |
| Fimbrial adherence determinants | Pef | pefA | 10 | Enteritidis |
| Fimbrial adherence determinants | Pef | pefB | 10 | Enteritidis |
| Fimbrial adherence determinants | Pef | pefC | 10 | Enteritidis |
| Fimbrial adherence determinants | Pef | pefD | 10 | Enteritidis |
| Serum resistance | Rck | rck | 10 | Enteritidis |
| Nonfimbrial adherence<br>determinants | ShdA | shdA | 10 | Enteritidis |
| Toxin | SpvB | spvB | 10 | Enteritidis |
| Secretion system | TTSS-2 translocated effectors | spvC | 10 | Enteritidis |
| Regulation | Regulates spvA, spvB and spvC | spvR | 10 | Enteritidis |
| Fimbrial adherence determinants | Lpf | lpfD | 11 | Enteritidis |
| Nonfimbrial adherence determinants | RatB | ratB | 11 | Enteritidis |
| Stress adaptation | SodCI | sodCI | 11 | Enteritidis |
| Iron uptake | Ent siderophore | entE | 38 | Kentucky |
| Adherence | K88 fimbriae | faeC | 38 | Kentucky |
| Adherence | K88 fimbriae | faeD | 38 | Kentucky |
| Adherence | K88 fimbriae | faeE | 38 | Kentucky |
| Iron uptake | Ent siderophore | fepC | 41 | Kentucky and Senftenberg |
| Disinfectant resistance/Iron uptake | SitABCD | *sitABCD* | 51 | Kentucky and Enteritidis |

Virulence factors, antimicrobial resistance genes and plasmid replicons were identified using the respective databases; Virulence Factor Database (via ABRICATE), Comprehensive Antimicrobial Resistance Database (via RGI) and the plasmidFinder database (via ABRICATE). Only genes which are unique to 2 or less serovars are listed.

### Serovar Kentucky and Senftenberg isolates harbored unique iron acquisition virulence factors

Serovar Kentucky isolates, regardless of the flock cohort they were obtained from, harbored similar virulence factor profiles (**Figure 3**). Unique to Kentucky isolates was the iron acquisition gene *entE* (involved in enterochelin synthesis) and shared with serovar Senftenberg isolates was the iron transport gene *fepC* (involved in enterochelin transport) (Bearson *et al*., 2008). Iron regulation is important for intracellular pathogens such as *Salmonella* as it is required for growth. For example, tight regulation of intracellular iron is crucial due to the detrimental effect of excess intracellular iron (Andrews, Robinson and Rodríguez-Quiñones, 2003). These iron regulation-related genes, in addition to those uniquely harbored by Kentucky and Senftenberg isolates, suggest an enhanced ability to regulate iron acquisition.

### Serovar Kentucky and Enteritidis isolates harbor distinct fimbrial adherence genes

Pathogenic bacteria employ a host of cell surface adhesins to adhere and eventually colonize novel host tissues such as the gasterointestinal tract (Kuijpers *et al*., 2019; Božić *et al*., 2020). Serovar Kentucky and Enteritidis isolates, each, harbored a distinct adhesion-related operon. Serovar Kentucky harbored the entire *fae* operon (*fae* of the K88 F4 fimbrial gene cluster as observed from our VFDB analysis) (**Table S1**).

Serovar Enteritidis isolates uniquely harbored 13 virulence factors which were present in a majority of Enteritidis isolates (**Figure 3**). These genes included the plasmid encoded fimbrial operon, pefABCD, responsible for adhesion to the intestinal epithelium (Seribelli *et al*., 2020) and 3 genes of the spvABCDR operon (*spvB, spvC,* and *spvR*) which is associated with survival and replication in macrophages (Rychlik, Gregorova and Hradecka, 2006). Lastly, the *rck* (resistance to complement killing) gene was identified in Enteritidis isolates and is known to increase serum resistance and adhesion to epithelial cell lines (Heffernan *et al*., 1994). These virulence genes aid in colonization and survival within host tissues and are similar to the functions of the unique VFs identified in Kentucky isolates. However, Kentucky isolates were isolated from the third flock cohort, while no Enteritidis isolates were isolates. These data suggest that the litter environment may impose a selective pressure on these virulence genes and the isolates that harbor them.

### *Salmonella* isolates harbored similar AMR profile

Forty-one antimicrobial resistance genes (ARGs) were identified using the Resistance Gene Identifier (RGI), which utilizes the Comprehensive Antibiotics Resistance database (CARD) (Alcock *et al*., 2020). Of these ARGs, 39 were present in 100% of the isolates (**Figure 4**). The 41 ARGs were predicted to confer resistance to 26 different antibiotic drug classes (**Table S4**). *kdpE* gene, a transcriptional activator of a two-component potassium transport system and *Bla*_TEM-60_ (identified as an hypothetical protein using Prokka and NCBI BLAST search), an extended-spectrum beta-lactamase gene, were identified in 38 and 40 isolates, respectively. *kdpE* has been reported to regulate virulence genes (Hughes *et al*., 2009; Zhao *et al*., 2010; Freeman, Dorus and Waterfield, 2013) and was found in all serovars from this study. *Bla*_TEM-60_ was present in all serovar Kentucky and Senftenberg isolates and absent from all Enteritidis isolates (**Table 3**).

**Figure 4.**
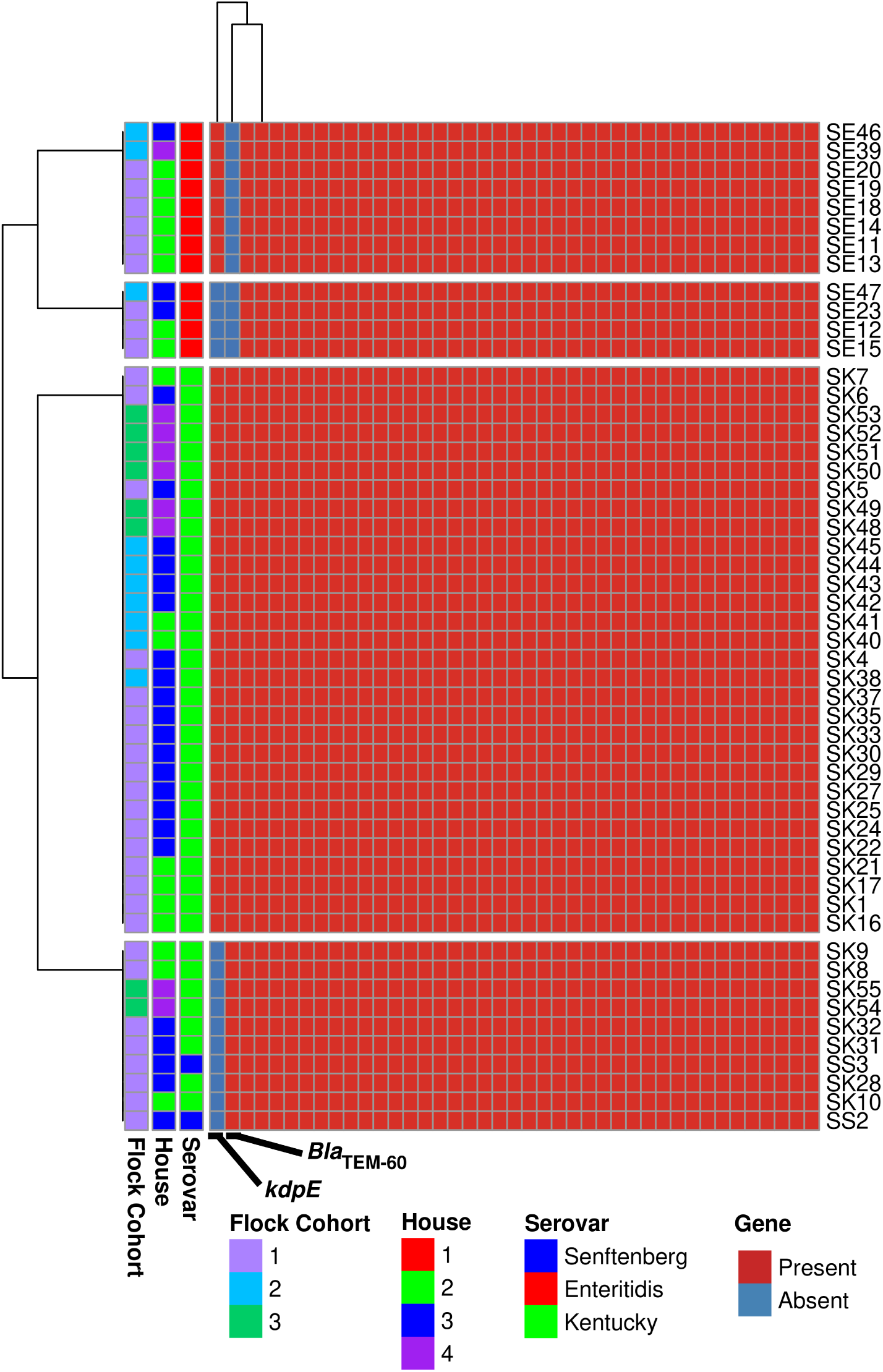
Heatmap of antimicrobial resistance genes identified. Antimicrobial resistance genes were identified from RGI and ResFinder. Heatmap was generated in R v4.0.4 with pheatmap v1.0.12 (clustering_method = “average” (UPGMA)) using the RGI and ResFinder data from **Table S1**.

Additionally, identification of acquired ARGs (genes likely to be harbored on a mobile genetic elements such as plasmids) via ResFinder (Bortolaia *et al*., 2020) resulted in only two hits; the *sitABCD* operon and the *aac(6’)-Iaa* gene. The *sitABCD* operon was present in all serovar Enteritidis and Kentucky isolates and absent from Senftenberg (**Table 3**). However, the *sitABCD* operon was at 72% percent identity which was below the 80% coverage identity cutoff for inclusion in our analysis. SitABCD is a putative iron transporter which plays a role in iron acquisition during infection and is important for growth in tissues following invasion of intestinal epithelium (Janakiraman and Slauch, 2000). The *aac(6’)-Iaa* gene, a chromosome-encoded aminoglycoside acetyltransferase (identified as *AAC(6’)-Iy* gene using the CARD database) was also present in all isolates (**Table S1**). *aac(6’)-type* genes have commonly been found in *S.* Kentucky and Typhimurium strains, however a study of 2,762 isolates harboring *aac(6’)-type* or *aac(6′)-Iy* gene found that only 11 isolates exhibited phenotypic resistance to aminoglycosides (Neuert *et al*., 2018)

In summary, the majority of the identified ARGs were chromosomally encoded and did not confer phenotypic resistance to the predicted antibiotics. This conclusion was supported by antibiotic susceptibility testing (AST) results. AST showed that 54/55 isolates were susceptible to the fourteen antibiotics tested (**Table S5**), while one isolate (SK32) exhibited resistance to streptomycin. We did not find any known ARG or chromosomal mutation that could explain the streptomycin resistance observed in SK32. Taken together, the *Salmonella* isolates in this study do not harbor ARGs capable of conferring clinically relevant levels of antibiotic resistance except *S*. Kentucky strain SK32.

### Co-occurrence of virulence factors and plasmid contigs

Five plasmid replicons were identified from the PlasmidFinder database (Carattoli *et al*., 2014) within serovar Kentucky isolates; “IncI1_1_Alpha”, “IncX1_3”, “Col156”, “Col(BS512) “and “Col(MG828)” (**Table 3, Table S6**). IncI1 and IncX1 were identified in all Kentucky isolates (38/38). A serovar Kentucky isolate (SK4) also harbored the Col plasmid replicons Col156 and Col(BS512) on short contigs of lengths 3,904 bp and 3,137 bp, respectively (**Table S6**). A total of 129 genes were identified on the IncI1 and IncX1 contigs, using the contigs from isolates SK1 and SK16, respectively (**Table S7**). The IncI1 contigs harbored silver and copper resistance operons (i.e., *silABCPRS* and *pcoABCESR,* respectively) as well as conjugation proteins (PilI, PilJ, PilK, PilL, PilM, PilN, PilO, PilP, TraA, TraB and TraC). A serovar Kentucky isolate was selected for long read sequencing to confirm the presence of the IncI1 plasmid within serovar Kentucky isolates. These results confirmed an 81,816 bp plasmid harboring both the *silABCPRS* and *pcoABCESR* operons (**Figure S1**).

IncFII(S), IncFIC(FII) and IncFIB(S) replicons were identified in serovar Enteritidis isolates (8/11, 8/11, 10/11 isolates, respectively) (**Table S6**). Rapid Annotation using Subsystem Technology (RAST) annotation of the isolate SE13 contig containing all 3 IncF replicons identified 97 genes (**Table S7**). This contig harbored the *spvABCDR* operon, *pefABCD* operon and *mig-5* genes in addition to the IncF plasmid conjugative proteins FinO, TraV, TrbD, TraA, TraB, TraK, TraE, TraL and TraA (**Figure S2**). No plasmid replicons identified in either *S.* Kentucky or *S.* Enteritidis isolates were lost over the multiple flock cohorts. These data suggest that VF and plasmid replicons remained constant within and across serovars. VFs and plasmid replicons harbored by *S.* Kentucky and *S.* Enteritidis isolates may be under specific environmental pressure from the litter environment.

## Discussion

The purpose of this study was to characterize the genomes of *Salmonella* isolates recovered from peanut hull-based broiler litter during the grow-out of 3 consecutive flocks of broiler chickens. Our objective was to identify ARGs, VFs and metal resistance genes harbored by the *Salmonella* isolates as well as understand how management and environmental factors can lead to genomic changes or persistence over multiple flock cohorts. Other studies, such as the one performed by Roll et al (Roll, Dai Prá and Roll, 2011) have demonstrated that *Salmonella* could persist over multiple broiler flock cohorts but no studies to the authors’ knowledge have characterized the genome of *Salmonella* isolates from flock to flock.

We isolated *S.* Kentucky, *S*. Enteritidis and *S.* Senftenberg serovars in the litter of the broiler houses studied (**Table 1**). Serovar Kentucky had the highest prevalence and persisted from flock 1 to 3. It is not surprising that *S.* Kentucky was the most dominant serovar in our study as it is one of the most commonly isolated poultry serovars from domestic chicken samples in the United States (Dunn *et al*., 2022). *S.* Kentucky was also the major serovar in breeder flocks from 2016 to 2020 (Siceloff, Waltman and Shariat, 2022). Unique to serovar Kentucky isolates were the VFs of the K88, F4, fimbrial *fae* operon and the iron uptake-associated gene *entE*. Also, unique to serovar Kentucky isolates were the IncX1and IncI1 plasmids, of which the IncI1 plasmid harbored copper and silver resistance operons in addition to conjugation related genes (**Table S7, Figure S1**). Kentucky isolates harbored multiple metal resistance operons which may have conferred a competitive advantage to surviving within the production environment, resulting in their persistence in the 3 flock cohorts. It is possible that the waterers and water lines acted as reservoirs for these isolates as copper sulfate was added to the drinking water. Addition of acidified copper sulfate is a common practice to sanitize water lines and waterers (Scott *et al*., 2018). However, while Senftenberg isolates harbored these operons on the chromosome, no Senftenberg isolates were observed during the second and final flock cohorts. It is plausible that metal resistance genes harbored on the chromosome poses a higher fitness cost on the bacterial host (e.g., *S*. Senftenberg) compared to when they are carried on plasmids (e.g., *S*. Kentucky). Our results suggest that *S*. Kentucky has evolved with VFs and plasmids which allow it to colonize broiler chickens and persist in the broiler house environment.

Virulence factor differences were explained by serovars (**Figure 3**), nevertheless all serovars harbored VFs with functions relating to secretion systems, fimbrial adherence determinants, invasion, iron uptake, macrophage inducible genes, magnesium uptake and nonfimbrial adhesion determinants (**Table S3**). *Salmonella* Enteritidis isolates harbored additional set of 13 VFs that were not present in either Kentucky or Senftenberg isolates. These VFs included plasmid genes (*spvABCDR* operon, *pefABCD* operon and *mig-5*) located on IncFIB and IncFII plasmid replicon contigs (**Figure S2**). These VFs have been identified in human cases associated with serovars Typhimurium and Enteritidis (Kuijpers *et al*., 2019; Seribelli *et al*., 2020). As these VFs encode for functions conferring increased fitness and survival within host tissues, is it possible that the chickens acted as a reservoir for these strains over multiple flock cohorts, and through the copraphagic nature of chickens, were continually deposited in and re-ingested from litter. Interestingly, house 4 had no detectable *S.* Enteritidis in the first flock however an Enteritidis isolate was obtained during the grow-out of the second flock. Similarly, no *S.* Kentucky isolates were obtained during the first 2 flocks in house 4 however during the third flock cohort 8 Kentucky isolates were obtained. This suggests that human or rodent transmission between houses and/or failed detection by the methods used are possible reasons for this observation (Backhans and Fellström, 2012). Additionally, it is possible that these isolates were introduced from the hatchery upon placement of the flock 3 cohort of chicks.

We found that while all *Salmonella* isolates harbored a core set of ARGs and VFs (95% and 72%, respectively) (**Figure 4, Figure 3**), each set of isolates grouped by serovar harbored a distinct set of ARGs, VFs, and plasmids (**Table 3**). While ARG identification resulted in 42 chromosome encoded ARGs, AST revealed that only one isolate was resistant to streptomycin (**Table S4**). This low prevalence of AMR is not surprising since the integrated farm adopted a “No Antibiotics Ever” program during grow-out (Oladeinde *et al*., 2023). Therefore, it is plausible that no antibiotic selective pressure existed in the litter.

We have provided new data on the genomic characteristics of *Salmonella* serovars Kentucky, Enteritidis and Senftenberg found in peanut hull-based litter during pre-harvest broiler chicken grow-out. We demonstrated that AMR was similar across all serovars while VF and plasmid profiles varied with respect to each serovar. Serovar Enteritidis harbored IncF plasmid associated replicons that were located on the same contigs as virulence factors associated with pathogenicity in humans, while serovar Kentucky isolates harbored an IncI1 plasmid harboring copper and silver resistance operons. While some *S.* Kentucky isolates persisted from flock 1 to 3, no *S.* Enteritidis isolates did. Therefore, these data suggest that some *Salmonella* serovars and strains are equipped with genetic factors that allow them to persist in broiler house environments under specific management practices including “NAE” programs that administer copper sulfate via drinking water. Nonetheless, there are several limitations of the study which could have biased our interpretation of the results including the small number of flocks monitored and the limit of detection of the method used for *Salmonella* isolation. Similarly, the initial broiler house cleaning procedures may have failed to remove all residual contaminants from previous flocks and subsequently resulted in these strains colonizing the litter environment. Thus, it is plausible the cleaning procedures resulted in differential results across each broiler house. Lastly, the results described were from one broiler farm that reused peanut hull-based litter and may not be representative of all farms that reuse litter for poultry production.

## Materials and Methods

Methods involving litter management on-farm, sample collection of litter as well as microbiological methods for bacterial isolation have been previously described (Oladeinde *et al*., 2023). Materials and methods used in this study, and others re-described from the previous work, is presented below.

### Study design

Three cohorts of broiler flocks were raised in succession in 4 co-located integrated commercial broiler houses in South Georgia between February and August 2018. Each of the broiler houses contained 22,000 to 24,000 broilers per flock. Each of the four co-located broiler houses underwent a full cleanout before fresh peanut hull-based litter placement. Each cohort was raised on the previous flock’s litter without any cleanout between and only mechanical conditioning was performed to remove caked portions. A commercial litter acidifier was applied during the downtime between flock cohorts for ammonium control (typically 1 week before sampling). Half house brooding was performed for the first 14 days of each flock. This practice restricted the birds to the front section of each broiler house. Additionally, copper sulfate was given via drinking water. Management practices and procedures employed are within the scope of routine industry practices.

### Litter sampling

Litter samples (n = 288) were collected across four broiler houses during early (chick age between 4-14 days old) and late grow-out (32-38 days old) for three consecutive flocks. This amounted to 6 sampling times per broiler house; each flock was sampled at two separate time points. Each house was divided into 12 subsections; four sections from front to back and each section was divided into three subsections from left to right (figure 1). During sampling, three litter grab samples were taken from each of the 12 subsections and pooled into one Whirl Pak bag. Pooled litter samples were then transported on ice to the United States National Poultry Research Center for further processing.

### *Salmonella* isolation and identification

All samples were processed within 24 h of collection. Thirty grams from each pooled litter grab sample was mixed with 120 ml phosphate buffered saline and shook with a hand wrist shaker (Boekel Scientific, Model 401000) for 10 min. Litter eluate (100 µl) was direct plated to both Brilliant Green Sulfur (BGS) and Xylose Lysine Tergitol-4 (XLT-4) agars. Plates were incubated 18-24 h at 37°C. Additionally, aliquots (1 ml) of the eluate were enriched in buffered peptone water (9 ml) for 18-24 h at 37°C. Enrichments were plated to BGS and XLT-4 agars and transferred to GN (Gram Negative) Hajna and Tetrathionate broths and incubated 24 h and 48 h, respectively at 37°C. Afterwards, 100 µl of GN Hajna and Tetrathionate broth Broths were then transferred to Rappaport-Vassiliadis R10 (RV) broth (BD; Franklin Lakes, NJ) and incubated at 37°C for 18-24 h. Thereafter, 10 µl of RV broth was plated to both BGS and XLT-4 agars. Isolated colonies characteristic of *Salmonella* were struck onto triple sugar iron and lysine iron agar slants and incubated at 37°C for 18-24 h. Presumptive *Salmonella* isolates (n = 55) were serogrouped with antisera (Becton Dickinson) and then cryopreserved.

### Antibiotic susceptibility testing

Antimicrobial susceptibility of *Salmonella* isolates was determined using the Sensititre™ semi-automated system (Thermo Fisher Scientific, Kansas City, KS) according to manufacturer’s instructions. Briefly, bacterial suspensions equivalent to a 0.5 McFarland standard were prepared, aliquoted into a CMV4AGNF panel and incubated at 37°C for 18 h. Minimum inhibitory concentrations were determined and categorized as resistant according to Clinical and Laboratory Standards Institute (CLSI) guidelines when available (CLSI, 2019); otherwise, breakpoints established by the National Antimicrobial Resistance Monitoring System (NARMS) were used (https://www.fda.gov/media/108180/download). Clinical and Laboratory Standards Institute. 2019. Performance standards for antimicrobial susceptibility testing, 30th ed. CLSI Document M100-Ed30. Clinical and Laboratory Standards Institute, Wayne, PA.

### Whole genome sequencing, processing and taxonomic classification

Select *Salmonella* isolates recovered from litter underwent Illumina short read sequencing and Oxford Nanopore long read sequencing. All DNA extraction was performed using the FastDNA spin kit for soil. Short read libraries were prepared using Nextera XT DNA library preparation kits (Illumina, Inc., San Diego, CA) following the manufacturers protocol. Libraries were sequenced on the MiSeq platform with 250-bp paired end reads. Four isolates underwent sequencing at Novogene (Novogene Co., Ltd., Tianjin) using the Illumina NovaSeq 6000 platform with 150-bp paired end reads. Long read sequencing was conducted at Novogene using the GridION platform (Oxford Nanopore Technology). Sequencing read quality control, adaptor and quality trimming, genome assembly, antimicrobial resistance gene identification, virulence factor identification, plasmid replicon identification, phage region identification and genome annotation were done using Reads2Resistome pipeline v0.0.2 (Woyda, Oladeinde and Abdo, 2023). Isolates selected for both short read and long read sequencing were assembled using both short and long reads using the --hybrid option in the Reads2Resistome pipeline. ResFinder (Bortolaia *et al*., 2020) was utilized for annotation of acquired resistance genes. Antimicrobial resistance gene identification was performed using the Resistance Gene Identifier v5.1.1 (RGI) which relies on the Comprehensive Antimicrobial Resistance Database and antibacterial biocide and metal resistance genes were identified with BacMet. MLST was determined using the mlst software (Seemann T, mlst Github https://github.com/tseemann/mlst), which utilizes the PubMLST website (https://pubmlst.org/). Serovar identification was performed using SISTR (Yoshida *et al*., 2016) via Reads2Resistome. The minimum identity match used for all reference database query hits was 85%. Verification of genes, as well as suspect plasmid replicon-containing contigs was done using Megablast implemented in Geneious Prime version 2022.2.2. *Escherichia coli* UMNK88 plasmid (Genbank accession CP002730.1) was used as a reference for the *fae* operon. Plasmid replicons were identified using the PlasmidFinder database via <u>ABRICATE</u> (Carattoli *et al*., 2014; Seemann, no date). Additional gene annotation, with available subsystem annotation, was performed using the Rapid Annotation using Subsystem Technology (RAST) (Aziz *et al*., 2008). Plasmid maps were generated using BLAST Ring Image Generator (Alikhan *et al*., 2011).

### Statistical analysis

Antimicrobial resistance genes identified through RGI were filtered to include hits which matched >=95% to the reference database sequence. Genes identified through BacMet and ABRICATE (virulence factors) were filtered to include those matching >=80% to the reference database. A table was generated based on the presence/absence of the identified genes in each isolate from each database. Correspondence analysis on the presence/absence table of antimicrobial resistance genes and virulence factors was conducted in R using the following packages: factoextra v1.0.7, FactoMineR v2.4 and corrplot v0.2-0. Heatmaps were generated using the pheatmap package in R. A distance matrix was generated using the jaccard metric via the vegdist function from the vegan v2.6-4 package (Jari Oksanen *et al*., 2022). The distance matrix was used to determine the optimal number of clusters and was implemented using the silhouette method from the fviz_nbclust function from the factoextra v1.0.7 package. Via the pheatmap function, hclust() from the stats v3.6.2 package was then utilized to perform hierarchical clustering using the ‘average’ (UPGMA) method. All analyses were done in R v4.0.4 utilizing RStudio v1.2.1106. Using the Illumina quality-controlled short-read sequences, taxonomic classification was performed using Kraken2.

## Data Availability

All raw isolate data, in FASTQ format, is available from NCBI under the accession number: **XYZ**. All other data are available upon request.

## Acknowledgments

We are grateful to Ms. Denice Cudnick, Dr. Kim Cook and Mr. Wilson for their logistical and technical assistance. This work was supported by the USDA Agricultural Research Service (Project Number: 6040-32000-012-000D). R.W. was supported under the National Institutes of Health (Grant number: 5T32GM132057-04 and 5T32GM132057-03). This research was partially supported in part by an appointment to the Agricultural Research Service (ARS) Research Participation Program administered by the Oak Ridge Institute for Science and Education (ORISE) through an interagency agreement between the U.S. Department of Energy (DOE) and the U.S. Department of Agriculture (USDA) (CRIS project number: 60-6040-6-009). ORISE is managed by ORAU under DOE contract number DE-SC0014664. All opinions expressed in this paper are the author’s and do not necessarily reflect the policies and views of USDA, DOE, or ORAU/ORISE. Any mention of products or trade names does not constitute recommendation for use. The authors declare no competing commercial interests in relation to the submitted work. USDA is an equal opportunity provider and employer.

## Supplemental Figures

**Figure S1.**
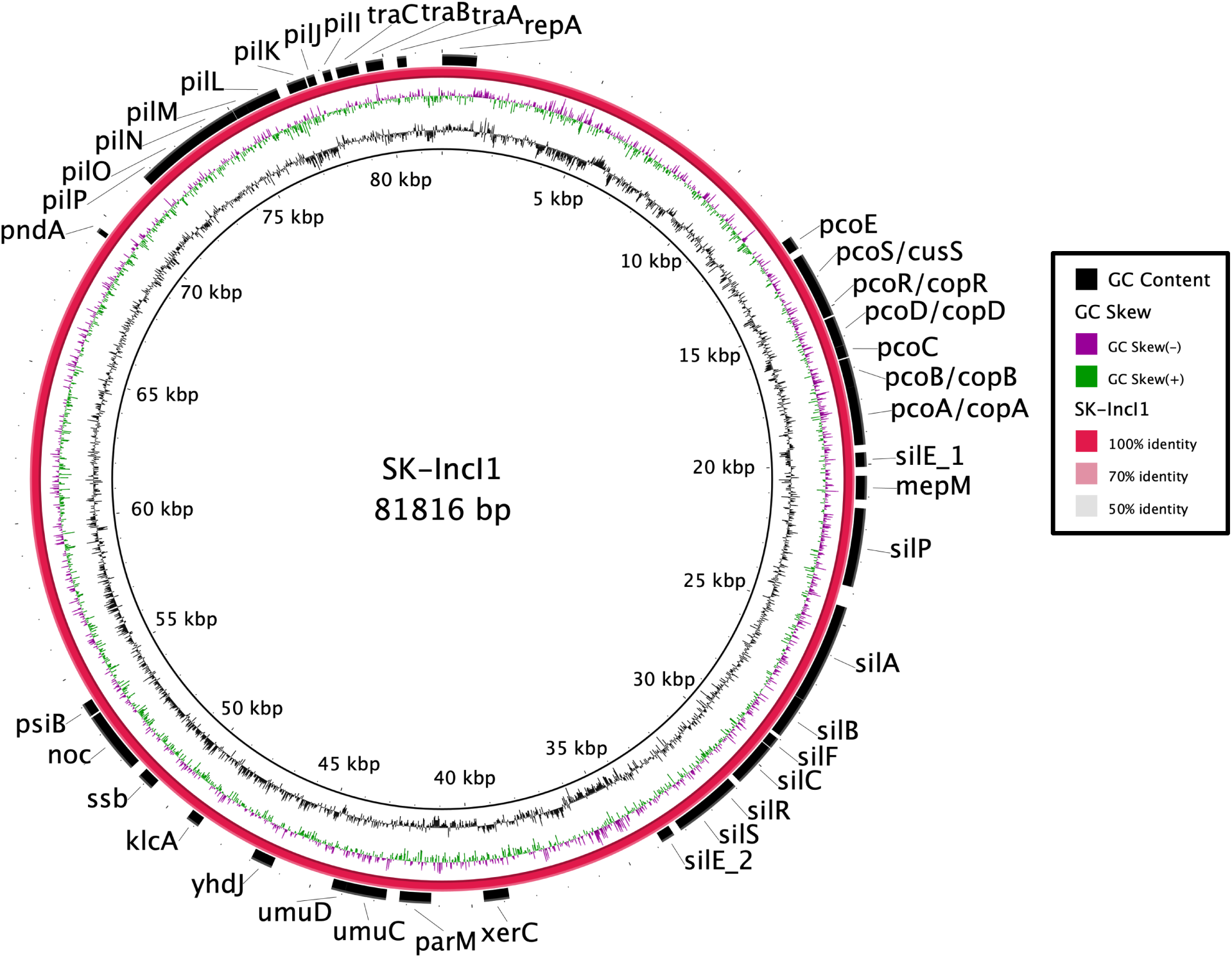
Annotated *Salmonella* Kentucky IncI1 plasmid.

**Figure S2.**
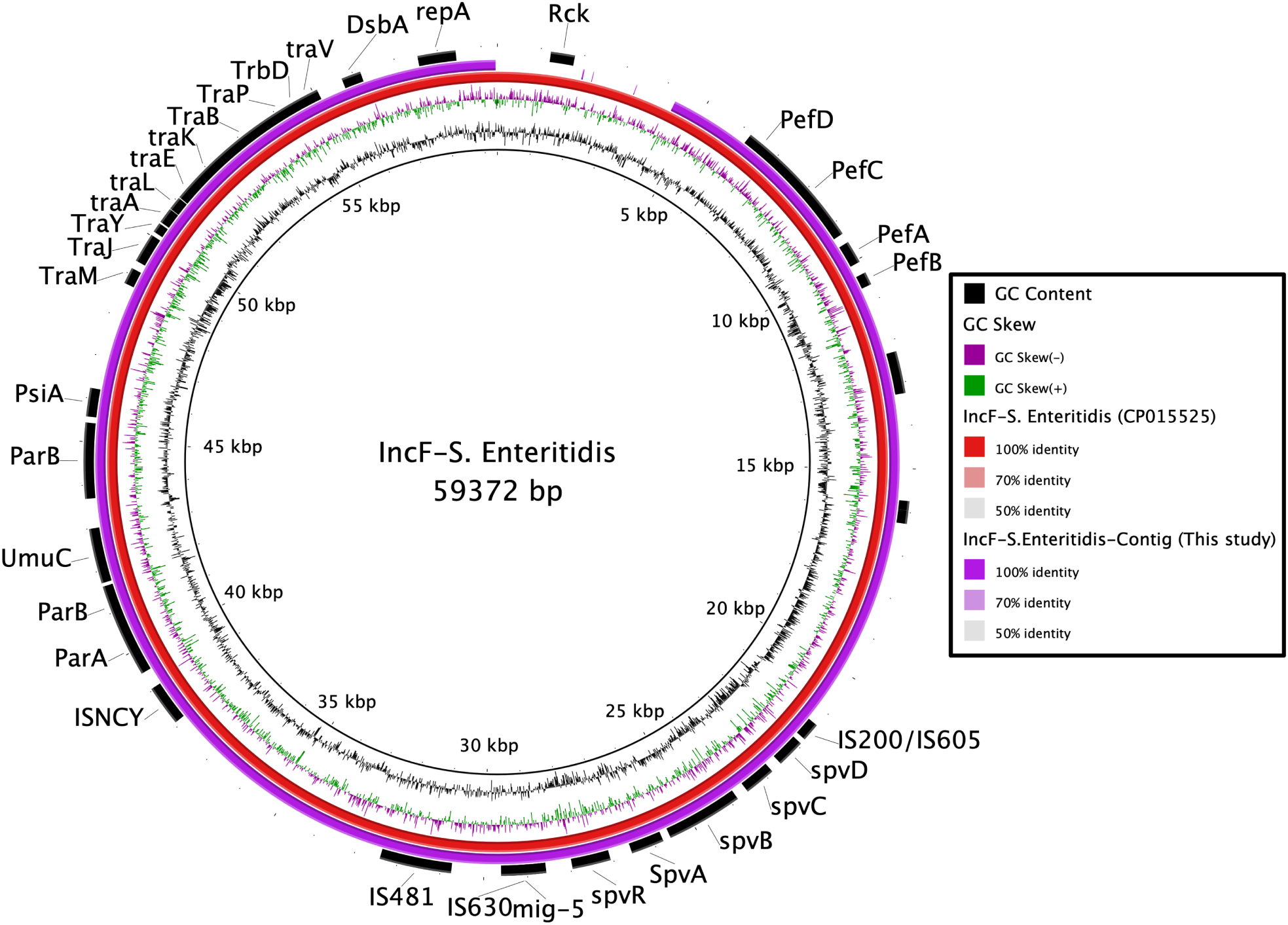
BLASTn alignment of the *Salmonella* Enteritidis IncF plasmid to the plasmid pSJTUF10978 (GenBank accession # CP015525).

## Supplemental Tables

**Table S1:** Metadata and ARG and VF presence absence spreadsheet (separate file)

**Table S2:** BacMet identified biocide and metal resistance genes (separate file)

**Table S3.** Virulence factor functional categories of virulence factors identified from the Virulence Factor Database (VFDB) (via ABRICATE (Reads2Resistome) (separate file)

**Table S4.** Drug classes conferred by genes identified by the Resistance Gene Identifier which utilizes the Comprehensive Antibiotic Resistance Database (separate file)

**Table S5.** Antibiotic susceptibility testing results for Salmonella isolates (separate file)

**Table S6.** *Salmonella* isolate-associated plasmid replicons (separate file)

**Table S7.** RAST plasmid annotations (separate file)

